# Gravitational expectations simultaneously attract and repel perception

**DOI:** 10.64898/2026.06.15.732061

**Authors:** Nick Simpson, Kirsten Rittershofer, Emma K. Ward, Matan Mazor, Clare Press

## Abstract

Perception is typically biased towards prior expectations. In some cases, however, it seems repelled away from expectations, such that percepts appear less like what is expected. Even more intriguingly, separate studies have recently reported that predictions derived from gravity may shape perception in opposing ways. Specifically, gravity causes unsupported objects to accelerate downwards, leading to two predictions; that objects will move downwards (location prior) and at an increasing speed (acceleration prior). There is evidence that perceptual judgements are attracted towards location priors yet repelled from acceleration ones. Here we examine these effects in the same paradigm to determine whether they result from different types of stimuli and judgement, or more interestingly, might result from opposite influences of common predictive mechanisms influencing perception. We first replicate previous reports of a systematic bias to report upward moving objects as more accelerating than downward moving objects: effectively a repulsion from acceleration priors. We then show that the effect applies both at the level of retinal space and due to contextual cues concerning gravitational direction. Finally, we find that participants’ errors in a location reproduction task are similarly consistent with a repulsion from acceleration priors and, simultaneously, with an attraction towards location priors. We conclude by considering the ways in which these concurrent attractive and repulsive biases may reflect mechanisms optimising fast, accurate, and informative experiences in our ever-changing sensory world, therefore optimising the interface between perception and learning.

**Public Significance Statement:** In a series of behavioural experiments, we show that expectations about how objects move due to gravity concurrently attract perception towards the prediction that objects move downwards, and repel perception away from the prediction that they do so at an increasing rate. These opposing influences inform current theories of perceptual processing, which explain how expectations may generate percepts that are fast, veridical, and informative.

## 1. Introduction

Gravity acts on objects such that they accelerate downwards at a rate of 9.8m/s^2^ (Isaac Newton, 1687). Animals should therefore possess an expectation in line with these statistics (Clark, 2013b, 2013a; Helmholtz, 2005; Mcintyre et al., 2001; Moscatelli & Lacquaniti, 2011) – either via hardwired processes or through learning about the statistical regularities in the environment (Fiser & Lengyel, 2026). Perhaps counterintuitively, however, people are biased to report downward moving objects as *less* accelerating than upward moving objects (Phan et al., 2024). This repulsion away from a gravitational ‘acceleration prior’ is counterintuitive because people are usually biased *towards*, not away from, perceiving what they expect (Bar et al., 2006; de Lange et al., 2018; Sterzer et al., 2010; Teufel et al., 2018; Yuille & Kersten, 2006) – a bias which is thought to render perception more accurate in a noisy sensory world.

Intriguingly, when asked about the position of unsupported objects, expectations about gravity yield the more commonly observed attraction biases: objects held up in the air are perceived as lower in the visual field than they are (De Sá Teixeira, 2016; Hubbard, 1997, 2020; Nagai et al., 2002). This attraction to a location prior – often termed ‘representational gravity’ – is also thought to be determined by gravitational expectations because it is mediated by the speed, mass and type of the objects (Hubbard, 1997, 2020; Nagai et al., 2002). Together, people thus seem to perceive unsupported objects closer to where they expect them due to gravity, but the changing speed of objects as less like what they expect due to the same gravitational expectations.

The co-occurrence of a repulsion from gravitational acceleration priors in tandem with an attraction towards location priors is potentially a critical datapoint for theorising about a so-called ‘perceptual prediction paradox’ (Press et al., 2020). Specifically, it is unclear why perceptual decisions are sometimes biased towards, and sometimes away from, expectations, when both biases seem like they should be adaptive. Attractive biases would optimise accuracy in a noisy sensory world, while repulsive biases would act against accuracy but would be more beneficial for learning - highlighting the deviations between expectation and reality. However, it might simply be that these two opposing effects are the result of different stimuli (e.g., real world scenes [Phan et al., 2024] vs artificial stimuli [Hubbard, 1997]), or of task requirements (a categorical report [Phan et al., 2024] vs continuous reproduction [Hubbard, 1997]). For example, while the attractive position effects seem more likely a result of perceptual processes because they have been shown across a number of different perceptual tasks (Hubbard, 2020), the repelling acceleration effect has only been shown in one paradigm and where decisional explanations are likely (Phan et al., 2024). The present studies therefore sought to understand whether the mechanism underlying acceleration prior repulsions is similar to, or overlapping with, that underlying location prior attractions. In so doing, we seek to understand how we optimise speed and accuracy of experience alongside informativeness (Press et al., 2020) — a question with a range of implications for perception, learning, and decision making.

Experiment 1 sought to replicate the recently reported effect of acceleration prior repulsion (Phan et al., 2024). Experiment 2 asked whether this repulsion was seen across two axes of potential influence – retinotopic and contextual sky-to-ground. These experiments thus determined whether there are robust and flexible influences of acceleration priors on direct reports about perception, which is arguably a crucial component of evidence concerning what we perceive. Experiment 3 used a location reproduction task similar to those used in the location prior literature to look for further evidence of acceleration repulsion in a paradigm where decisional interpretations are less likely (Sánchez-Fuenzalida et al., 2023). This final manipulation importantly allowed us to also measure the effect of acceleration and location priors in the same task, using the same stimuli, and in the exact same experimental trials. Convergence across such distinct tasks would increase the likelihood that these priors are indeed influencing perception, rather than explained via other biases that are hard to dissociate in single tasks.

To anticipate our findings, across all three studies we consistently find evidence of perceptual repulsion from gravitational acceleration priors. Experiments 1 and 2 show that people report downward moving objects as less accelerating than upward moving objects, across both retinotopic and contextual axes. In Experiment 3, we find that a repulsion from gravitational acceleration priors exists alongside an attraction towards location priors. We therefore consider how these opposing effects may reflect common mechanisms optimising perception to generate fast, accurate, and informative experiences in our ever-changing sensory world.

## 2. Methods and Results

### 2.1 Experiment 1

#### 2.1.1 Motivation

Experiment 1 (preregistered: https://doi.org/10.17605/OSF.IO/3MW9S) aimed to replicate the repulsion from acceleration priors found by Phan et al. (2024). Participants were presented with upward and downward moving basketballs that accelerate, decelerate, or travel at constant velocity on different trials. They were required to report whether the ball was accelerating or decelerating. We thus sought to determine whether they are biased to report downward moving objects as more decelerating than upward moving objects.

#### 2.1.2 Participants

We calculated that a sample size of 100 participants would give an approximate statistical power of 98% to find an effect size of 0.4 - as observed by Phan et al. (2024). We recruited 135 participants via the Prolific platform with normal or corrected to normal vision, and they completed the experiment online. Participants were removed from analysis if their overall accuracy in the task was below 75%, leaving 100 included participants. After data collection, and in deviation from our pre-registration, we removed participants with any point of subjective constant velocity outside the maximum range (see below for details), leaving 85 analysed participants (mean age = 40.1 years; sd = 12.7). The experiment lasted ~15 minutes and participants were paid a small honorarium for their time. The study was approved by the local ethics committee and participants gave informed consent before starting the experiment.

#### 2.1.3 Procedure

On each trial, a basketball court was displayed over 80% of the screen height and a basketball travelled vertically through the centre of the court (Figure 1.A). The ball could be accelerating, decelerating, or travelling at a constant velocity. Once the ball had left the screen, participants pressed the ‘a’ key if they thought it was accelerating and the ‘d’ key if they thought it was decelerating. There was no time limit for responses and trials started 1500 ms after the previous response.

**Figure 1.**
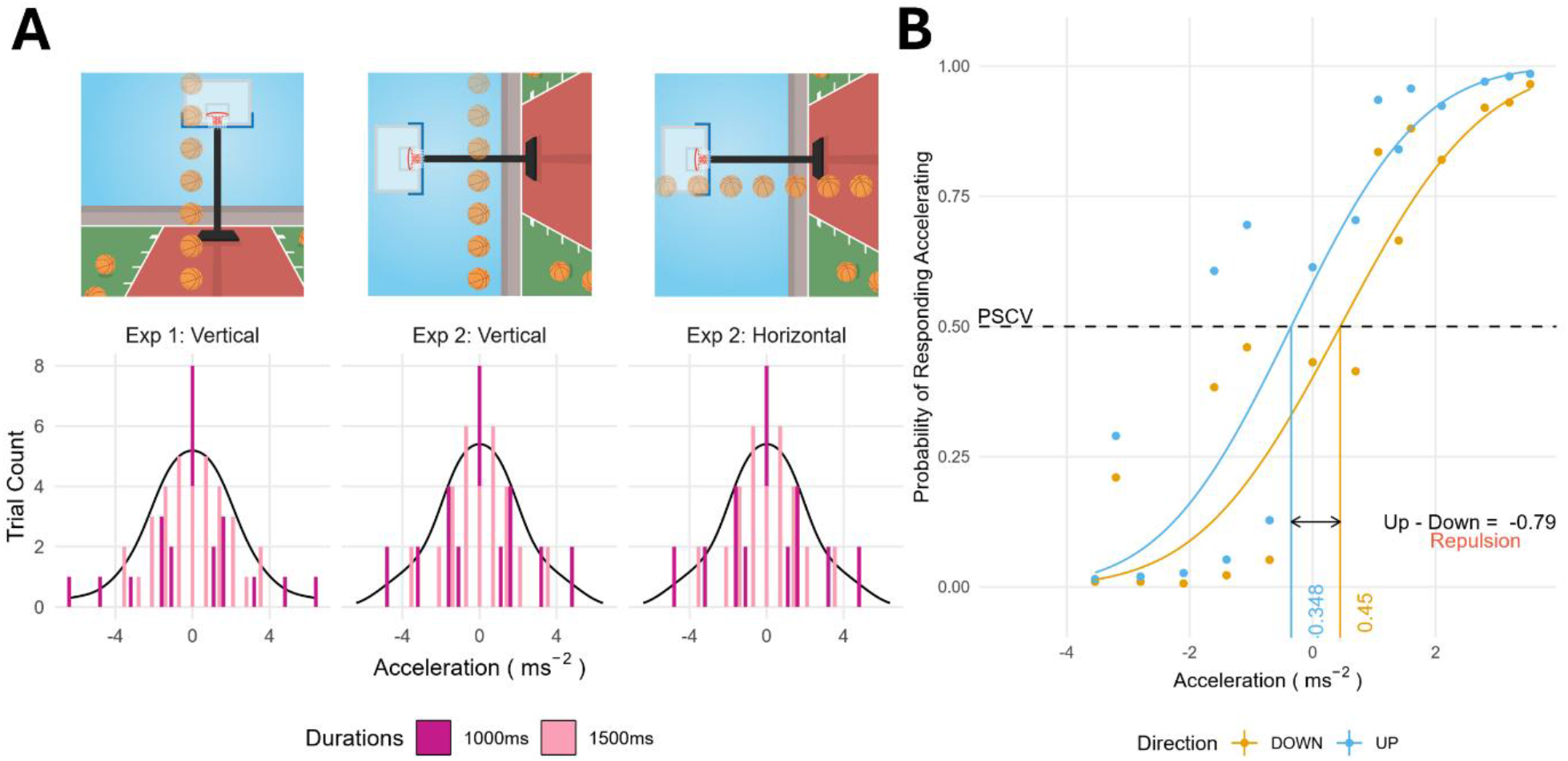
Stimuli. **A** In Experiment 1, participants saw a basketball court displayed over 80% of their screen height. A basketball travelled across the screen either upwards or downwards. Each motion direction was presented with the same acceleration values centred around constant velocity. The histogram bar colours represent the time for which the ball was on the screen. Participants were asked to respond whether they thought the ball was accelerating or decelerating. In Experiment 2, the same basketball scene was presented but rotated by 90 degrees. Therefore, when balls travelled up and down, contextual gravity equivalently influenced up and down motion, but retinal expectations differed. In contrast, balls travelling left and right appeared to be moving up and down within the contextual display, but not in retinal coordinates. As a result, effects observed in the horizontal plane would be generated by contextual, rather than retinal, expectations. Experiment 2 acceleration distributions were identical for horizontally and vertically moving balls. **B** An example psychometric curve fit to participant responses. Separate curves were fit for up and down trials, and a PSCV value extracted as the point at which participants were equally likely to respond accelerating or decelerating. A more positive PSCV for downward trials compared to upward trials shows a repulsion from gravitational expectations.

Following the procedure of Phan et al. (2024), and to avoid the start or end velocity being perfectly correlated with acceleration, the ball remained on the screen for one of two screen durations: 1000 ms or 1500 ms. For each screen duration, 11 equally spaced acceleration values were chosen from the range 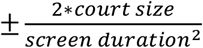. An approximately Gaussian distribution of acceleration values was presented to each participant for both upward and downward moving balls (Figure 1a Exp 1). The contextual cues on the screen (e.g. basketball hoop) were designed such that the height and width of the display would correspond to approximately 4m in the real world. As the ball appeared and disappeared at the sides of the screen, the motion calculations used a total distance of 4.4 m. This size gave an acceleration range of ±8*ms*^−2^, which aimed to approximate naturally occurring values.

We presented 8 blocks of 25 trials, resulting in 100 trials for each condition (i.e., upward and downward moving balls). The trial order was pseudo-randomised using the Mersenne Twister pseudorandom number generator, initialised in a way that ensures registration time-locking (Mazor et al., 2019). Vertical trials were alternated with horizontal trials, which will be analysed in a future publication along with other manipulations presented in the horizontal.

Prior to the main task, participants were given detailed instructions, followed by 16 practice trials where accuracy feedback was given.

#### 2.1.4 Results and Discussion

The point of subjective constant velocity (PSCV) was calculated for each participant and motion direction (up/down). The PSCV was extracted from psychometric curves as the point at which the participant was equally likely to respond accelerating or decelerating (Figure 1B). Psychometric curves were fit using the quickpsy R package with acceleration value as the explanatory variable and participant, screen duration, and direction as grouping factors (Linares & López-Moliner, 2016). Curves were fit only to the preregistered acceleration values in the range ±3.6*ms*^−2^, and participants were removed who had a PSCV value outside the range of ±8*ms*^−2^ (i.e., outside the range of presented stimuli), leaving 85 participants. The mean PSCV across screen duration was calculated to give PSCV values for each direction and participant. A more negative PSCV value represents a bias to report the motion as more accelerating, relative to decelerating.

Upward motion had a more negative PSCV than downward motion (*t*(84) = −6.80, *P* < 0.001; Figure 2.A and B Exp 1), representing a bias to report upward motion as more accelerating than downward motion. The bias effect was observed when controlling for the slope of the psychometric curves and was corroborated by a model free analysis (see Supplementary Materials 1 and 3).

**Figure 2.**
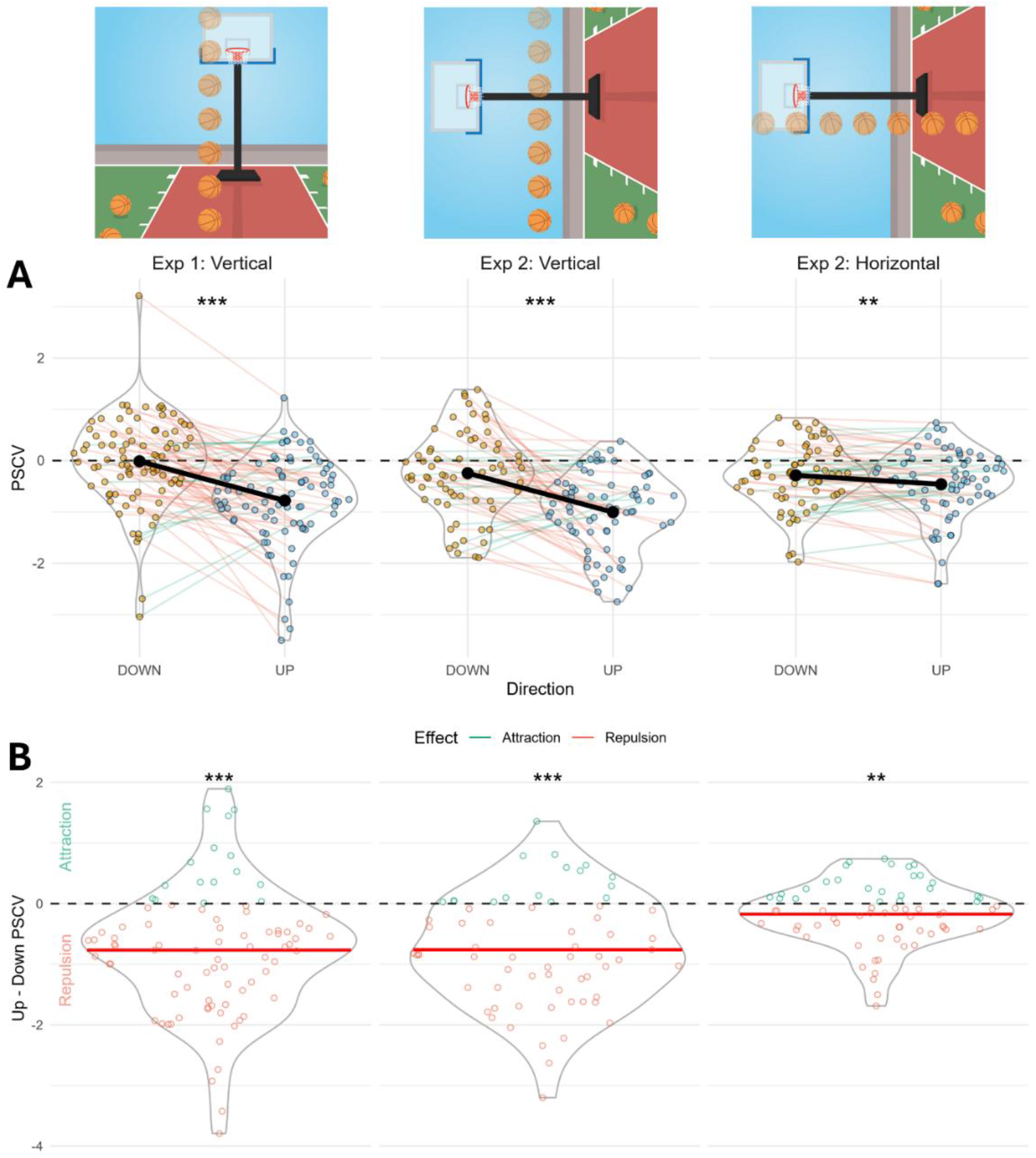
Experiment 1 and 2 Results. **A** Extracted PSCVs compared across conditions for each Experiment. Coloured points represent individual participant PSCV values and black points show the mean across participants. The dotted black line shows a PSCV of zero. A more negative PSCV value for upward compared to downward motion, across both retinal and contextual axes, represents a repulsion from gravitational expectations. **B** The difference between the PSCV values for upward and downward trials shown in A. Coloured points represent individual participant differences (PSCV for UP minus PSCV for DOWN) and the red line shows the mean difference across participants. The negative mean value demonstrates the repulsion from gravitational expectations.

The results from Experiment 1 are therefore in line with Phan et al. (2024), whereby upward motion is reported as more accelerating than downward motion. These results suggest that perceptual decisions are biased away from gravitational acceleration priors that free-falling objects accelerate downwards, and projected objects decelerate upwards.

### 2.2 Experiment 2

#### 2.2.1 Motivation

The effect observed in Experiment 1 likely reflects repulsion from a gravitational acceleration prior. Experiment 2 (preregistered: https://doi.org/10.17605/OSF.IO/D895R) sought to determine whether this bias is yoked to retinal or contextual space. If the effect is only present in retinal space, it may be due to in-built properties of the visual system, such as interactions between hemifield sensitivity (Levine & McAnany, 2005). If it remains in contextual space, then it is more likely that there is a component of the effect that relies upon more flexible prediction processes which can reflect interactions between learning and perception. To decouple the retinal and contextual expectations, we presented participants with both vertically and horizontally moving balls on a basketball court rotated by 90 degrees. In this way, balls travelling along the horizontal were unaffected by the retinal contributions of gravitational expectations, but the contextual direction of motion differed between leftward and rightward balls. Conversely, vertically moving balls were unaffected by contextual direction but the retinal contributions of gravitational expectations differed between upward and downward balls.

#### 2.2.2 Participants

Participants were recruited via the Prolific platform and completed the experiment online. This experiment used a Bayesian stopping rule to complete data collection when a Bayes Factor exceeded 3 or fell below 1/3. Data were collected from 99 participants. Thirty-one participants were removed from analysis because their overall accuracy in the task was below 75%. Additionally, participants were removed from analysis if any PSCV was outside the tested range, leaving 67 participants (mean age = 35.6 years; sd = 11.7) in the horizontal condition and 66 participants (mean age = 35.2 years; sd = 11.5) in the vertical condition. The experiment lasted around 15 minutes and participants were paid a small honorarium for their time.

#### 2.2.3 Procedure

The procedure for Experiment 2 was largely identical to that in Experiment 1, with two differences. First, in this experiment the basketball court was rotated by 90 degrees such that contextual gravity acted horizontally, and retinal gravity acted vertically. Second, balls travelled both horizontally and vertically.

We presented 8 blocks of 28 trials, following the identical distributions in Figure 1.A Exp 2: Vertical and Horizontal. The background rotation alternated every block between +90 and −90 degrees such that the contextual gravity acted equally often in each direction. The distribution was altered subtly from Experiment 1 to allow for identical trials to travel in both rotated contexts (Figure 1.A). The trials were pseudo-randomised, ensuring registration time-locking (Mazor et al., 2019).

#### 2.2.4 Results and Discussion

The analysis for Experiment 2 was identical to that for Experiment 1, but PSCVs were calculated now for four conditions: retinal up (vertical motion), retinal down (vertical motion), contextual up (horizontal motion), and contextual down (horizontal motion). Additionally, following the preregistration, participants were rejected if any of their PSCV values fell outside of the tested range of ±4.8*m*/*s*^2^. Our preregistered analysis used Bayesian statistics (leading to the same conclusions and reported in Supplementary Materials 2) but for consistency we report the frequentist statistics here. To compare the retinal and contextual direction expectation effects, we performed a two-way repeated measures ANOVA with PSCV as the dependent variable and plane (horizontal or vertical) and direction (up or down) as predictors. This analysis found a significant main effect of both direction (*F*(1,64) = 53.66, *P* < 0.001) and plane (*F*(1,64) = 13.91, *P* < 0.001), and a plane by direction interaction (*F*(1,64) = 21.24, *P* < 0.001). This interaction was driven by a repulsion regardless of plane, but a larger repulsion in the vertical (*t*(65) = −6.81, *P* < 0.001; Figure 2.B Exp 2 Vertical) than horizontal axis (*t*(66) = −2.78, *P* < 0.01; Figure 2.A and B Exp 2 Horizontal).

These findings therefore suggest that both retinal and contextual gravitational expectations contribute to repulsion effects, albeit with larger retinal contributions^1^.

### 2.3 Experiment 3

#### 2.3.1 Motivation

Experiments 1 and 2 demonstrated that gravitational expectations produce a repulsion effect in a two-alternative forced choice (2AFC) task in both contextual and retinal dimensions, suggesting the effects may reflect more flexible prediction processes which perhaps mediate interactions between learning and perception. This repulsion effect was observed, however, in a setting where participants were asked to make explicit binary judgments about the acceleration of objects, leaving open the possibility that the observed repulsion effects reflect biases in post-perceptual, decisional processes (Gregory, 1980; Ratcliff et al., 2016; see also Firestone & Scholl, 2015). It is possible, for example, that because most people are aware that objects should accelerate downwards due to gravity, they believe that they will be biased to see such acceleration. As such, they may compensate for this bias at a decisional level when explicitly judging acceleration. If the compensation is larger than the underlying perceptual bias, repulsion will emerge.

In Experiment 3 (preregistered: https://doi.org/10.17605/OSF.IO/6TRCX; https://doi.org/10.17605/OSF.IO/8YWCV), we therefore asked whether acceleration repulsion can be observed in a task where decisional biases are less likely, and indeed one where attraction towards location priors has been demonstrated (Hubbard, 2020). Specifically, we asked participants to reproduce the position of a flash of a moving ball, noting that reproduction tasks have been argued to minimise such decisional contributions relative to perceptual effects (Sánchez-Fuenzalida et al., 2023)^2^.

#### 2.3.2 Participants

Three data sets were collected via the Prolific platform with the same paradigm – an exploratory sample of 79 participants (mean age = 36.4 years; sd = 11.1), a follow-up pre-registered sample of 150 participants (mean age = 38.6 years; sd = 11.5), and a second pre-registered replication sample of 500 participants (mean age = 35.1 years; sd = 8.17). Participants were removed from analysis if the correlation between their responses and the true flash location was below 0.3. This correlation value was chosen, and preregistered, based upon the exploratory data and the understanding that responses should be broadly in line with flash location, with a medium to strong effect. The experiment lasted ~20 minutes and participants were paid a small honorarium for their time.

#### 2.3.3 Procedure

Similarly to Experiments 1 and 2, participants were presented with a ball travelling vertically on the screen with a range of acceleration values. The acceleration distributions were also similar, with 100 trials for each direction (up/down), but all stimuli now lasted for 1000 ms (Figure 3B). In Experiment 3, the ball was now black and changed colour to dark grey for 50 ms at some point in the trajectory (Figure 3A). Once the ball had left the screen, the participant’s mouse location was presented as a small black circle which could only move vertically along the ball’s motion path. Participants were asked to click where they saw the ball flash.

**Figure 3.**
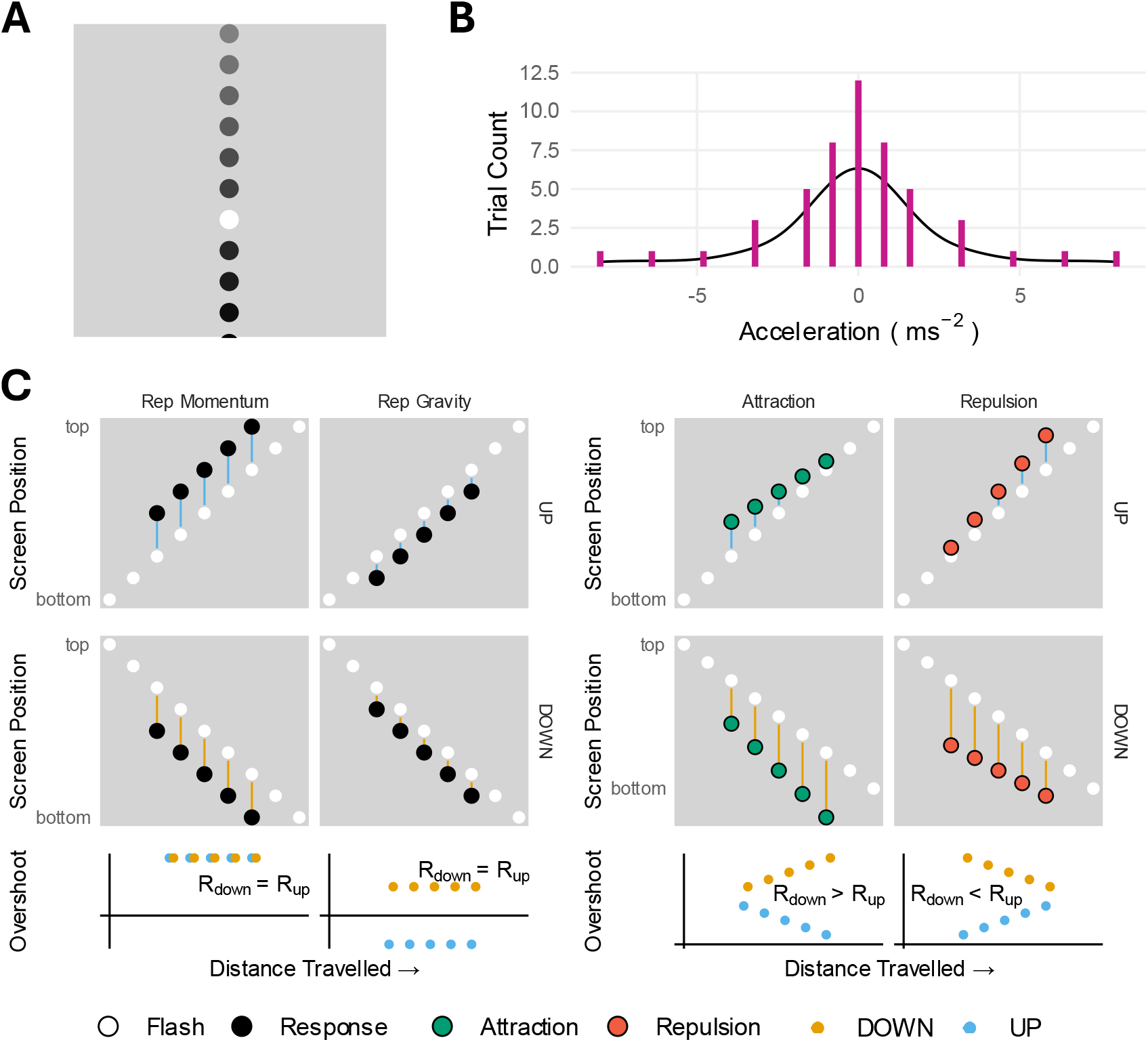
**A** Screen design for Experiment 3. Black balls travelled vertically and flashed a different colour along the trajectory. Participants were asked to click where the ball flashed after the ball left the screen. The white flash is exaggerated for demonstration and the transparency represents the ball’s downward motion. **B** Acceleration distribution for trials in Experiment 3. Identical acceleration values were presented for upward and downward trials. **C** Theoretical results from acceleration expectations impacting participants’ overshoots for constant velocity trials. Columns show different effects on location perception. The top row shows upward motion with the position on the screen on the y axis and the distance the ball has travelled on the x axis, the middle row shows downward motion, and the bottom row shows the overshoots from the upward and downward motion as a function of distance travelled. Left hand side: When travelling upwards, representational momentum and representational gravity (location prior) act in opposing directions and are constant across a constant velocity trial. They act in the same direction for downward motion. These two effects lead to no difference in overshoot across distance travelled (bottom row). Right hand side: Objects perceived to travel faster will have a larger overshoot because representational momentum increases with speed. Therefore, an attractive bias would predict larger overshoots at the bottom of the screen compared to the top of the screen. This results in a more positive correlation between the distance travelled and the overshoot for downward compared to upward trials (*R*_*down*_ > *R*_*up*_; bottom row). The opposite is true for a repulsion effect.

Different from the previous experiments, the background was presented as a light grey square rather than a basketball court. This change was implemented such that the visual information concerning the flash would not differ depending upon the background colour at the flash location, and also because Experiment 2 suggested that the contextual information is not necessary to generate the repulsive effect. The ball was also made half the size such that the flash locations were easier to specify. Finally, we presented the ball at the top or bottom of the screen for 1000 ms prior to the beginning of motion, to allow early flashes to be detected without reorienting.

The flash location was pseudo-randomised on each trial between 20% and 80% of the screen height. However, given that mis-localisations are limited by the edges of the screen, as pre-registered, we only analysed the trials that flash between 25% and 75% of the screen height. Prior to the main task, participants were given detailed instructions and 8 practice trials with non-directional accuracy feedback. The main experiment consisted of 10 blocks of 20 trials.

#### 2.3.4 Analysis

We first calculated the distance between the centre of the flash and the location clicked by the participant, given as a proportion of the screen height. The overshoot was positive if the participant reported the flash location as ahead in the trajectory, and negative if it lags.

The overshoot of a moving ball is often studied in paradigms whereby participants report the point of disappearance of a moving ball as further along its trajectory (Reed & Vinson, 1996). This ‘representational momentum’ is modulated by velocity, such that faster velocities generate more representational momentum (Freyd & Finke, 1986). Due to gravitational expectations, upward moving objects are expected to decelerate and thus to travel more slowly at later timepoints compared to earlier ones. Given an attractive bias, participants should therefore have *larger* overshoots early in the trajectory, with the reverse for downward moving objects (Figure 3C). Such an attraction effect would hence manifest as larger overshoots at the bottom of the screen where objects are expected to be faster. If the bias is repulsive, the opposite pattern would be predicted, such that overshoots are larger at the top of the screen where objects are expected to be slow.

To compare between upward and downward motion directions, we calculated the correlation between the size of the overshoot and the distance from the start of the motion at which the flash occurred, to see how the perception of velocity changes across the trajectory. An attractive bias would lead to a more positive correlation for downward trials compared to upward trials, and a repulsive bias would lead to the opposite. We compared the correlation coefficients at the group level with a paired samples t-test to understand whether there was a difference in the correlations for upward and downward moving trials.

#### 2.3.5 Results and Discussion

Figure 4A shows that participants mostly performed the task well, with all responding within a few ball widths of the centre of the flash. Consistent with representational momentum, participants reproduced the location of the flash with a mean overshoot of 1.71 ball widths (*BW*) in the Exploration sample (*t*(78) = 19.32, *P* < 0.001), 1.69 *BW* in the Replication sample (*t*(149) = 24.89, *P* < 0.001), and 1.56 *BW* in the Second Replication (*t*(499) = 37.81, *P* < 0.001).

**Figure 4.**
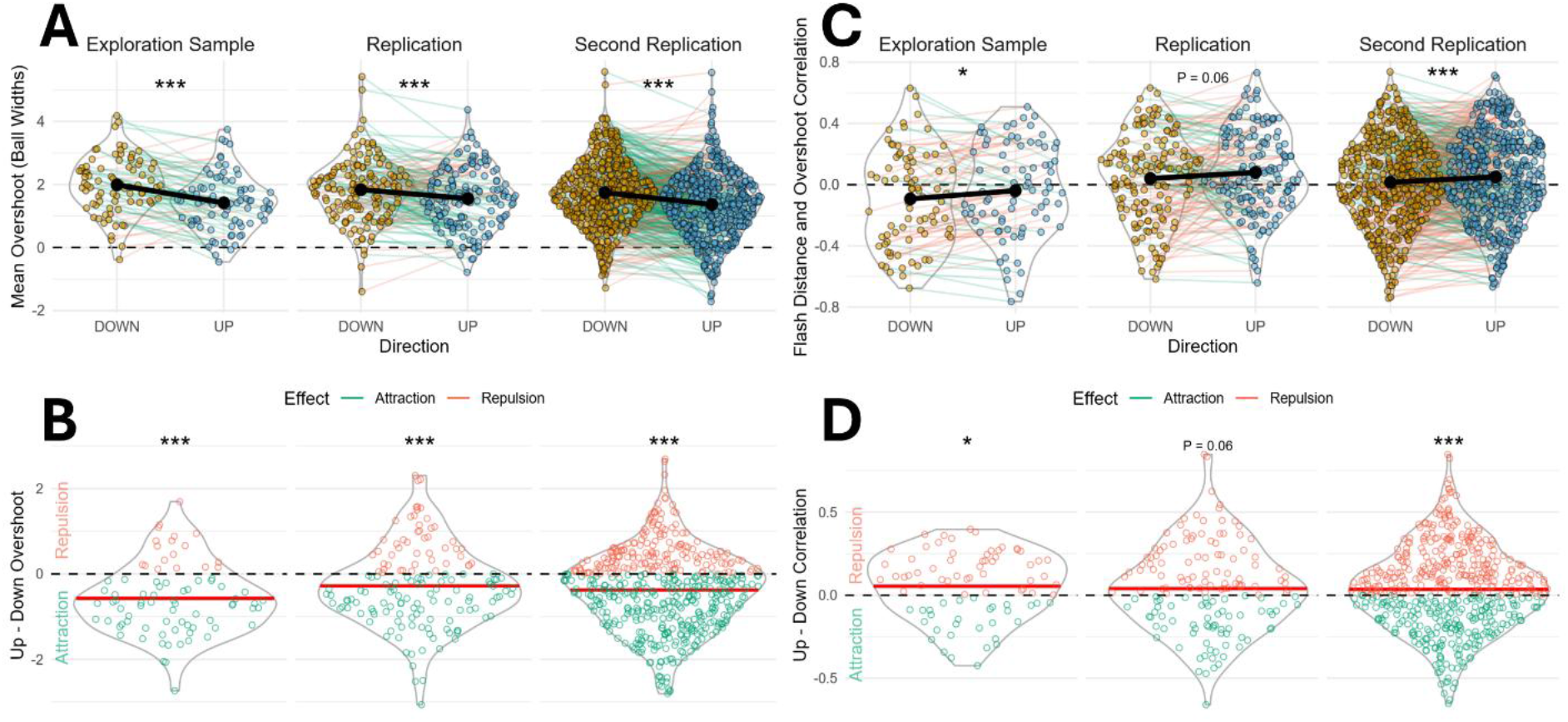
Experiment 3 results. **A** Participant mean overshoots separated for upward and downward trials. Coloured points show individual participants’ mean overshoot and the black points show the group average. **B** The difference in mean overshoots for upward and downward trials from A. Coloured points represent individual participant differences and the red line shows the group average. A negative difference demonstrates that upward trials have a smaller overshoot than downward trials. **C** The correlation between flash distance and overshoot. Coloured points show individual participants’ correlation coefficients and the black points show the group average. A more negative correlation for downward trials compared to upward trials indicates that downward trials are perceived as less accelerating - a repulsion from gravitational expectations concerning acceleration. **D** The difference in correlation coefficients for upward and downward trials from C. Coloured points show individual participant differences and the red line shows the group average. A positive difference shows a repulsive effect.

Supporting an attraction towards location priors, participants had a smaller overshoot for upward moving trials compared to downward moving trials (Figure 4A and B) in the Exploration sample (*Mean*_*UP*_ = 1.42 *BW, Mean*_*DOWN*_ = 1.99 *BW, t*(78) = −6.17, *P* < 0.001), the Replication (*Mean*_*UP*_ = 1.55 *BW, Mean*_*DOWN*_ = 1.83 *BW, t*(149) = −3.72, *P* < 0.001) and the Second Replication (*Mean*_*UP*_ = 1.37 *BW, Mean*_*DOWN*_ = 1.74 *BW, t*(499) = −9.30, *P* < 0.001). That is, location priors gave rise to an attractive bias of ~0.1-0.2 BW.

Figure 4C and D show the comparison of overshoot-flash distance correlations for upward and downward trials. In the Exploration sample, the correlation coefficient for upward trials was more positive than the correlation coefficient for downward trials (*t*(78) = 2.40, *P* < 0.05). In the Replication sample, the same trend existed but the two-tailed t-test was not significant (*t*(149) = 1.89, *P* = 0.060). In the Second Replication, the pre-registered one-tailed t-test replicated the Exploratory finding (*t*(499) = 3.18, *P* < 0.001). Moreover, in all three samples the effect met the significance threshold in a significant portion of window sizes other than the preregistered one (see Supplementary Materials 4).

The representational momentum, attraction towards location priors and repulsion from acceleration priors, can all be included in a linear model to test whether the additional complexity of all three effects together best describes the overshoot data. When comparing AIC values in a step-wise model comparison across the data from all experiments, the winning model included factors associated with all three effects (Eq. 1; See Supplementary Materials 6 for full analysis). Namely, the positive estimate of the effect of ball speed on overshoot shows the representational momentum effect (*β* = 2.246 × 10^−3^; *P* < 0.001), the negative estimate of the effect of direction on overshoot shows the attractive location prior effect (*β* = −9.357 × 10^−3^; *P* < 0.001), and the interaction between direction and distance travelled shows the repulsive acceleration prior effect (*β* = 2.208 × 10^−3^; *P* < 0.001).

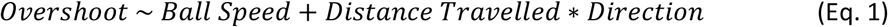

This combination of findings thereby broadly shows that perceptual judgements are also biased away from gravitational expectations of acceleration in a reproduction task. Namely, there is less of an increase in overshoot along the trajectory for downward trials compared to upward trials, indicating that downward motion is perceived to accelerate less than upward motion. This experiment, across three samples, thus replicates the finding of a repulsion from gravitational expectations about acceleration from Experiments 1 and 2 in a task where decisional explanations are less likely, and finds this effect alongside typical attractive location-prior effects.

## 3. General Discussion

The present studies sought to understand the nature of an intriguing influence of expectation on perceptual decisions, whereby predictions derived from gravity appear to shape perception in opposing ways. Gravity causes objects to accelerate downwards, leading to predictions that objects will move downwards (location prior) and that they will do so at an increasing speed (acceleration prior). There is evidence that perceptual judgements are attracted towards location priors yet repelled from acceleration ones, but these findings had been derived across different studies, and the acceleration repulsion has been observed only once and in a paradigm where decisional explanations are likely.

In Experiment 1, we first provide a conceptual replication of the recent demonstration of acceleration prior repulsion (Phan et al., 2024). Experiment 2 separates the retinal and contextual contributions to this effect by rotating the scene-based context by 90 degrees, finding contributions of both, albeit with a stronger contribution from retinal coordinates. Experiment 3 provides evidence for a similar repulsion when participants reproduce a location rather than making decisions directly about acceleration. The persistence of the repulsion across tasks likely demonstrates that these repulsive effects are driven by perceptual, rather than decisional, processes (Sánchez-Fuenzalida et al., 2023). Experiment 3 also supports previous findings that object location perception is biased downwards in line with the force of gravity (Hubbard, 2020). These experiments demonstrate that perception of objects in motion is concurrently attracted toward gravitational expectations of location and repelled from gravitational expectations of acceleration.

The observed repulsion effects are at odds with most findings in the predictive perception literature, whereby perception is usually attracted *toward* expectations (e.g. de Lange et al., 2018; Press et al., 2020). Why, therefore, would responses be biased away from gravitational acceleration priors when so often we are instead biased toward these expectations? Such a bias would render perceptual decisions less accurate, on average, given that objects more often accelerate downwards (Press et al., 2020). It may explain why we often rely on heuristics to make perceptual decisions about gravitational motion (Vicovaro et al., 2019), and could likely hinder our ability to interact with unsupported objects (Zago & Lacquaniti, 2005).

In fact, perceptual repulsion may also provide adaptive benefits, because it likely results from the amplification of the surprising features of our environment (Brown et al., 2013; Wolpert et al., 2003). It is often the surprising features that are more behaviourally relevant; we must react appropriately to unexpected events, and they sometimes reflect that we need to update our current world models (Mathys et al., 2011; Rao & Ballard, 1999). One might argue that in the case of gravity, we should rarely, if at all, update expectations concerning how gravity acts on free-falling objects, but we may instead update expectations concerning the forces acting on an object – e.g. reflecting animate forces or air resistance (Jörges & López-Moliner, 2017). Specifically, although motion against gravity is abnormal for a free-falling object, animate motion can produce a wide range of motion trajectories including upward acceleration (de Vries & Wurm, 2023; Nguyen & van Buren, 2023). Hence, an error signal highlighting deviation from gravitational motion could arguably lead us to switch to a prior that animate forces are at play (Gyulai, 2004; New et al., 2007; Szego & Rutherford, 2007; Tremoulet & Feldman, 2000).

Despite the rarity of expectation based perceptual repulsion, it may relate to some other phenomena in the literature. For example, repeated events generate a reduced neural response (Krekelberg et al., 2006; Rankin et al., 2009), which is thought to repel percepts from the previous event (Crane, 1988; Fritsche et al., 2017; Ward et al., 2024). Some have argued that such repetition phenomena are mediated partially by prediction mechanisms, but the relative contribution with respect to dissociable adaptation processes is a matter of debate (Feuerriegel, 2024; Feuerriegel et al., 2021; Summerfield et al., 2008; Todorovic et al., 2011). There are other types of judgement where unexpected events have been shown to be recalled better (Spaak et al., 2022), perceived with greater intensity at particular time ranges (Yon & Press, 2017), and sometimes perceived more readily (Nguyen & van Buren, 2023). Despite different types of effect, these may in principle be generated by similar mechanisms to facilitate our model-updating in cases of surprise.

There are a number of differences between the situations in which repulsive and attractive influences are found that may, in principle, drive the opposite patterns. The number of possible explanations is dramatically reduced by demonstrating both attractive and repulsive effects via the same prior and in the same paradigm, as in the present study. For example, even if we do not know whether gravitational expectations emerge ontogenetically or phylogenetically, a difference in origin for attracting and repelling priors, e.g., as proposed by Kilteni et al. (2019), becomes less likely with these patterns. It also seems unlikely that there is a different contribution of perceptual and decisional processes to the location- and acceleration-prior effects, given they are demonstrated in the same study and with the same task, potentially ruling out arguments for different effect directions for perceptual and post-perceptual processes (Fritsche et al., 2017; Moon & Kwon, 2022).

One exciting possibility is that the *precision* of the learned prior and incoming sensory information may determine the ultimate influence on perception. The opposing process theory (OPT) is a recent account which aims to explain these distinct influences of expectation on perception (Press et al., 2020). It postulates that perception is primarily biased towards expectations, to optimise the rapid generation of largely veridical percepts. However, if a stimulus generates an especially large error signal with respect to predictions (a large Kullback-Leibler Divergence [KLD] between prior and posterior), to the extent that it is unlikely reflective of sensory noise, a reactive process enhances perception of these model-breaking events, or perhaps elements, to optimise accurate model-updating. A large KLD would result from either the modes of the distributions being highly distinct or the distributions being highly precise. As gravity is a well-defined and ubiquitous prior (Jörges & López-Moliner, 2017), the precision would be high relative to those more commonly examined in statistical learning paradigms. Thus, this high precision may be why we observe repulsion effects here for acceleration, unlike in the majority of the predictive perception literature – especially noting that the other common repulsive effect in the literature, the size-weight illusion, could be explained similarly (Buckingham et al., 2009; Murray et al., 1999). Under this account, perception of location may be distinctly attracted downwards because at each particular timepoint during the motion there is little deviation between the expectation and the input, yet acceleration estimates require integration over multiple timeframes. Thus, the cumulative error signal is larger perhaps for acceleration. This cumulatively larger signal could cross the threshold to initiate the secondary process described by the OPT. However, to determine whether these repulsion effects are likely driven by such processes, future work could employ time-resolved examinations (see Rittershofer et al. (2026)) and manipulations of relative KLD across trials.

A different theory of perception grounded in efficient coding has recently been used to explain both attractive and repulsive effects (Wei & Stocker, 2015). The theory proposes that there can be differences in the precision of neural encoding along the continuum of certain stimuli and that a Bayesian observer will be repelled away from regions of high-precision whilst also being attracted to the prior (Hahn & Wei, 2024). The foundational phenomenon for this theory is that our perception of orientation is repelled from the cardinal axes and this can be explained by, for example, a higher density of cells coding for the cardinal orientations (Furmanski & Engel, 2000; Li et al., 2003). Under efficient coding accounts, a net repulsion would emerge when the rate of change of encoding resources (precision) across the stimulus space is higher than that of the prior (Hahn & Wei, 2024). For an efficient coding explanation of the current data, a high level expectation of gravitational motion (constant downward acceleration of 9.8*ms*^−2^) would thus need to govern the encoding resources and a short term expectation of the acceleration profiles in the experiment itself (wide range of expected acceleration values) would govern the prior. This would allow for the higher rate of change of encoding resources across the acceleration space than the prior. However, this explanation requires that the encoding system is able to change as a function of contextual expectations because the direction of motion determines the expected change of speed. Additionally, the mechanism of this dynamic encoding must be different from the mechanism of the attractive influence of the prior or the two influences would be indistinguishable. Despite this explanatory hurdles, efficient coding has been used to explain similar contextually dependent expectations (Bays, 2024; Fritsche et al., 2020; Prat-Carrabin & Woodford, 2022). Similar to the OPT, the efficient coding framework relies on differences in prior precision but additionally takes into account differences in neural resources and sensory and stimulus noise, thus – dependent upon particular assumptions – also providing an account of perception which may explain why gravitational expectations lead to both attractive and repulsive effects.

Motion perception has long been studied with respect to visual illusions such as the flash-lag effect, whereby a stationary object flashed next to a moving object is reported as offset in the opposite direction of motion (Nijhawan, 1994). EEG decoding evidence suggests that this type of phenomenon is due to an automatic motion extrapolation mechanism which reduces neural lag so that we are able to perceive moving objects in real time (Blom et al., 2020; Hogendoorn, 2020; Johnson et al., 2023). This same mechanism can explain the representational momentum effect, and potentially why objects are reported as lower than they truly were. However, it is less clear how these repelling effects of acceleration expectation would be supported by such mechanisms. Future work could establish the extent of neural lag to determine its modulation by gravitational expectations, and thus elucidate how our visual system simultaneously represents position and motion effectively (Blom et al., 2020; Johnson et al., 2023).

In conclusion, we find across three studies that perception is repelled away from gravitational expectations that objects *accelerate* downwards (acceleration priors), yet attracted towards gravitational expectations that objects *move* downwards (location priors). This repulsion effect is observed across perceptual paradigms, and across retinal space and contextual cues, suggestive of a robust and flexible phenomenon. This rare observation of expectation-based perceptual repulsion, especially in the context of perceptual attraction on other axes, may reflect common mechanisms optimising perception to generate fast, accurate, and informative experiences in our ever-changing sensory world.

## Constraints Against Generality

The studies reported here were conducted online with participants who have access to a computer and the internet, and the ability to read English and follow instructions during a difficult, though short-duration, experimental task. Participants were recruited regardless of their country of origin or their location at the time of data collection, but it is likely that we have recruited a sample that has higher than average education and access to resources. It is unclear how these results would generalise to the wider global population, and any claims about fundamental human cognition should be verified in more globally-representative samples.

## Transparency and Openness

We report how we determined our sample size, all data exclusions, all manipulations and all measures in the study. All data, analysis code and stimulus presentation code is available at. We pre-registered the initial design and hypotheses for this set of studies before data was collected – Experiment 1 https://doi.org/10.17605/OSF.IO/3MW9S, 2 https://doi.org/10.17605/OSF.IO/D895R, and 3 https://doi.org/10.17605/OSF.IO/6TRCX, https://doi.org/10.17605/OSF.IO/8YWCV.

## CRediT Roles

Nick Simpson: Conceptualization, Data curation, Formal analysis, Investigation, Methodology, Project administration, Software, Validation, Visualization, Writing – original draft, Writing – review & editing Kirsten Rittershofer: Conceptualization, Methodology, Software, Writing – review & editing Emma Ward: Conceptualization, Methodology, Software, Writing – review & editing Matan Mazor: Conceptualization, Investigation, Methodology, Supervision, Validation, Writing – review & editing Clare Press: Conceptualization, Funding acquisition, Methodology, Project administration, Resources, Supervision, Writing – review & editing

## Acknowledgements

This work was funded by a European Research Council (ERC) consolidator grant (101001592) under the European Union’s Horizon 2020 research and innovation programme, and Leverhulme Trust project grant (RPG-2022-358), both awarded to CP. We are grateful to other past and present members of the Action and Perception Lab for useful input throughout the studies.

## Footnotes

1 Interestingly, a previous study sought to distinguish retinal and contextual coordinates to gravitational influences on perception and found no influence of the latter. Miwa et al. (2019) tested whether body-line (retinal), contextual, and true (vestibular) directions of implied gravity affect the reported acceleration. When participants (N=7) were seated upright they presented upright and inverted contexts and found no contextual effects. However, when lying on their back, a 90 degree rotated context produced similar repulsion effects as described in Experiment 2. Our findings from a larger sample suggest that this previous study may have been underpowered to observe these smaller contextual contributions, relative to the larger retinal influences.

2 A similar study has been conducted to ask this question specifically in the context of acceleration priors, but we did not replicate their results (See Supplementary Materials 5) (Hubbard, 2001).

