## Supplementary Materials for "Gravitational expectations simultaneously attract and repel perception"

### Gravitational expectations simultaneously attract and repel perception: Supplementary Materials

#### 1. Preregistered Analyses Experiment 1

##### *1.1 Model Free Bias Analysis*

To supplement the psychometric curve analysis, a model free bias analysis was performed on all comparisons. The bias value was calculated by coding responses as 1 for accelerating and 0 for decelerating. For each participant, a mean response was found for each acceleration value, then the mean across acceleration values was found to give two screen duration dependent bias values for each condition. A single bias value was found by averaging across the screen durations.

Participants were more likely to respond accelerating for upward, than downward, moving balls ($t(99) = 10.12, P < 0.001$; Supplementary Figure 1 Exp 1). This analysis supports the PSCV analysis in the Main Text, and indicates that perceptual decisions are repelled from the expectation that objects accelerate downwards.

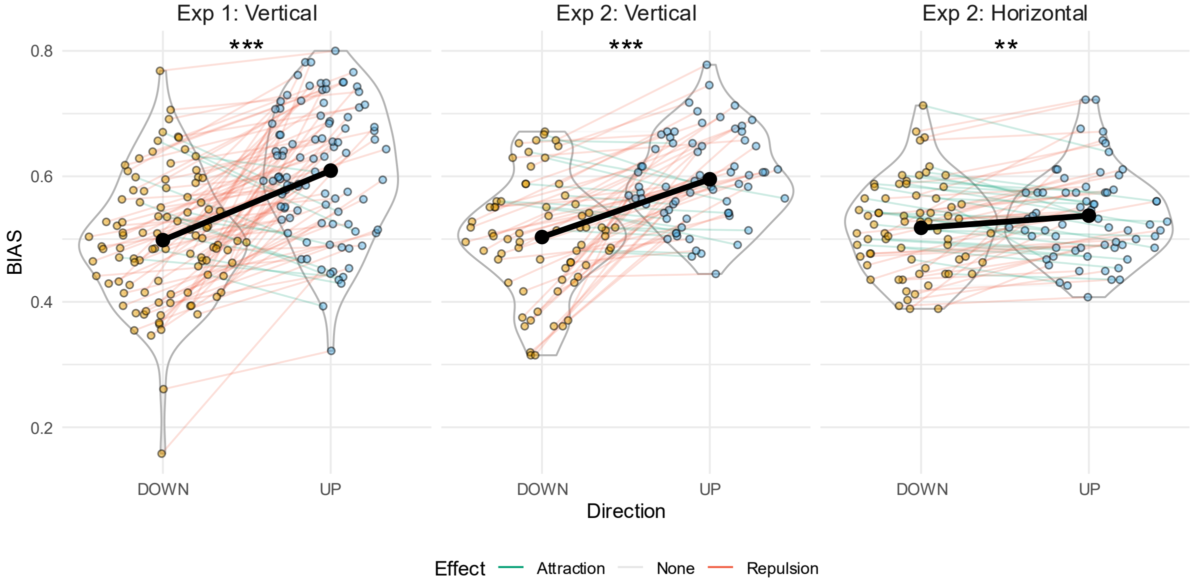


Supplementary Figure 1. Model free bias analyses. Coloured points refer to individual participants' response bias, where a larger value indicates more of a bias to respond 'accelerating'. The black points show the group mean. A larger bias value for UP compared to DOWN replicates the main text findings that upward motion is reported as more accelerating than downward motion.

*1.2 Initial/Final Speed Analysis*

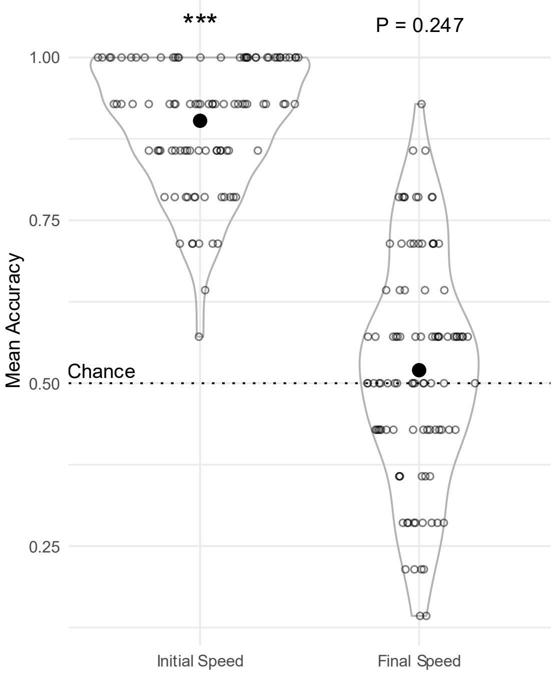


Supplementary Figure 2. Accuracy in trials where the initial or final speed is abnormal for the acceleration value. Hollow points show individual participants' mean accuracies and the filled points show the group average. The dotted line shows chance accuracy.

Across experiments, participants report whether the ball on the screen was accelerating or decelerating. If the ball were to remain on screen for a single duration, the initial speed for accelerating balls would always be lower than that for decelerating balls, and the final speed for accelerating balls would always be higher than that for decelerating balls. Therefore, we used two screen durations in these studies to reduce the informativeness of the start and end speeds, despite noting that this control is incomplete and therefore that future work must properly unpack the different contributors.

To test whether participants were exclusively using the start or end speeds, we compared the trials where the start/end speeds were atypical for their condition. There were two acceleration values where the start speeds were atypical – a deceleration of $-0.7ms^{-2}$ had an initial speed of $3.46ms^{-1}$ and a screen duration of 1500ms, whereas an acceleration of $1.067ms^{-2}$ had a higher initial speed of $3.87ms^{-1}$ because it had a screen duration of 1000ms. Therefore, accuracy on these trials required ignoring the start velocity value as an indication of acceleration value. Furthermore, for the final speed, an acceleration of $0.7ms^{-2}$ had a final speed of $3.46ms^{-1}$, whereas a deceleration of $-1.067ms^{-2}$ had a final speed of $3.87ms^{-1}$. By ensuring the accuracy of these specific trials is above chance, we can reduce the likelihood of the start or end speed being exclusively used for acceleration judgements. This analysis is only performed for the included participants.

The mean accuracy of trials with an acceleration value of $-0.7ms^{-2}$ and $1.067ms^{-2}$ was 90%, which is above chance ($t(99) = 41.51, P < 0.001$). Therefore, it is unlikely that the start speed of the balls was being used exclusively to judge acceleration in this study. The mean accuracy of trials with an acceleration value of $0.7ms^{-2}$ and $-1.067ms^{-2}$ was 52%, which is not different to chance ($t(99) = 1.16, P = 0.247$; see Supplementary Figure 2). We therefore cannot rule out that the final speed was contributing to judgements of acceleration perception in Experiment 1. The directional bias in reported accelerations is the matter of interest, and whether participants make this judgement based on the true change of speed, the final speed, or a combination of both, is arguably less central to our questions, but would be interesting to unpack further in future work.

This analysis was only pre-registered for Experiment 1. However, Experiment 2 Vertical ($MeanAcc_{Init} = 0.89$; $MeanAcc_{Fin} = 0.56$) and Horizontal ($MeanAcc_{Init} = 0.94$; $MeanAcc_{Fin} = 0.58$) trials showed a similar pattern of results, but both initial and final speed accuracy were now significantly above chance (Vertical: $P_{Init} < 0.001, P_{Fin} < 0.05$); Horizontal:($P_{Init} < 0.001, P_{Fin} < 0.01$). Therefore, it is easier to rule out that participants were entirely judging acceleration in Experiment 2 on final speed.

#### 2. Preregistered Analyses Experiment 2

*2.1 Model Free Bias Analysis*

See Section 1.1 for methods.

As Experiment 2 used a Bayesian stopping rule, Bayesian tests were preregistered, but for continuity with other studies we report the frequentist statistics in the Main Text. We here report the Bayesian tests, in line with the preregistration, and where the patterns were identical to those revealed by the frequentist statistics. Specifically, vertical comparisons show a significant difference between UP and DOWN biases ($t(67) = 8.16, P < 0.001; BF = 7.25 \times 10^8$; Supplementary Figure 1 Exp 2 Vertical). Similarly, the horizontal trials of Experiment 2 show a difference between contextual UP and DOWN biases ($t(67) = 2.80, P < 0.01; BF = 4.74$; Supplementary Figure 1 Exp 2 Horizontal).

To further complement the Main Text analysis, we used a 2x2 ANOVA with direction (up/down) and plane of motion (vertical/horizontal) to test whether the directional bias difference interacted with plane of motion. We again replicate the findings with the frequentist statistics. We find a main effect of direction ($F(1,67) = 70.46, P < 0.001; BF = 2.08 \times 10^4$) and plane of motion ($F(1,67) = 9.70, P < 0.01; BF = 3.72$), as well as an interaction between the direction and plane ($F(1,67) = 30.29, P < 0.001; BF = 26.81$).

*2.2 PSCV Analysis*

Similarly, the PSCV analysis patterns were identical with Bayesian statistics. There was a difference between upward and downward PSCVs for the vertical ($t(65) = -6.81, P < 0.001; BF = 3.09 \times 10^6$), and horizontal contextual upward and downward PSCVs ($t(66) = -2.78, P < 0.01; BF = 4.58$). Using a 2x2 ANOVA with direction (up/down) and plane of motion (vertical/horizontal), we find a main effect of direction ($F(1,64) = 53.66, P < 0.001; BF = 2.06 \times 10^4$) and plane of motion ($F(1,64) = 13.91, P < 0.001; BF = 3.72$), alongside an interaction between the direction and plane ($F(1,64) = 21.24, P < 0.001; BF = 27.30$ ).

#### 3. Slope Analysis Experiment 1 and 2

To analyse the precision of the fitted psychometric curves, the beta values (standard deviation of the cumulative normal distribution) for the slopes were extracted. Participants were excluded if they had a PSCV value outside the tested acceleration range and/or a beta value outside $\pm 3$ standard deviations of the mean. In Experiment 1, there was no difference in beta values for downward and upward trials ($t(85) = -1.86, P = 0.067$; Supplementary Figure 3 Exp 1: Vertical). In Experiment 2, participant beta values also showed no main effect of motion direction ($F(1,60) = 0.13, P = 0.723$) but did show a main effect of motion plane ($F(1,60) = 8.80, P < 0.01$). There was no interaction between motion direction and plane ($F(1,60) = 3.04, P = 0.086$). Pairwise comparisons of motion directions in Experiment 2 showed no difference between vertical up and down beta values ($t(62) = -0.76, P = 0.453$; Supplementary Figure 3 Exp 2: Vertical) or horizontal up and down beta values ($t(63) = 1.29, P = 0.203$; Supplementary Figure 3 Exp 2: Horizontal). Pairwise comparison of vertical and horizontal beta values showed that vertical beta values were larger than horizontal beta values ($t(120) = -2.89, P < 0.01$). To sum, there were no slope effects likely to generate our bias effects of interest.

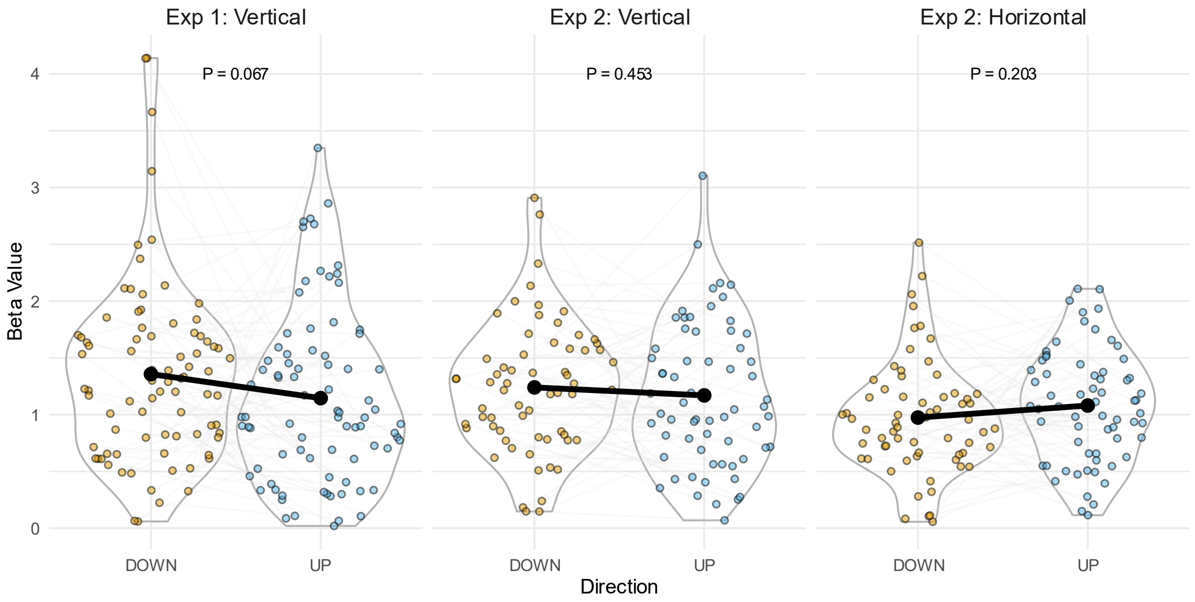


Supplementary Figure 3. Beta values (standard deviation of the cumulative normal distribution) for participant psychometric curve fits. Coloured points represent individual participant beta values and black points show the mean. Paired samples t-tests show no difference between beta values for upward and downward trials.

Nevertheless, in order to understand whether the PSCV effects were dependent on the slope values, the PSCV was modelled using linear mixed-effects models with direction and beta values as fixed effects and a random intercept for each participant (Eq. 1). By including the beta values as a covariate, we were able to estimate the unique contribution of direction on the PSCV. Experiment 1 showed that upward motion predicted a more negative PSCV value than downward motion ($\beta = -0.877, P < 0.001$) and larger beta values also predicted a more negative PSCV ($\beta = -0.275, P < 0.001$). Experiment 2 Vertical also showed the same direction effect ($\beta = -0.808, P < 0.001$) and beta effect ($\beta = -0.339, P < 0.01$). Experiment 2 Horizontal shows the same direction effect ($\beta = -0.125, P < 0.05$) but no beta effect ($\beta = -0.118, P = 0.185$). Thus, direction is a robust predictor of PSCV when accounting for participant beta values, and we can be confident that the observed bias effects are not due to slope differences.

$$lmer(PSCV \sim beta\ value\ +\ direction + (1\,|\,participant)) \qquad \text{(Eq. 1)}$$

**4. Pre-registered Window Analysis Experiment 3**

Participants were of course unable to report flashes as occurring off the screen. This dependency could artificially reduce biases when analysing flashes that occurred very early or late in the motion trajectory. However, we did not want to introduce an expectation that flashes occur in the centre of the screen as this could cause participants to ignore the majority of the motion trajectory. As such, we presented flashes at all locations but selected a window in which to analyse the data from 25% to 75% of the trajectory.

This selected window size was taken from the exploration sample (Supplementary Figure 4. Exploration Sample; A Window Size of 0.25 corresponds to data from 25% to 75% of the screen). However, this window size in the replication sample did not generate a significant effect while smaller windows did (Supplementary Figure 4. Replication). In both samples, the direction of effect is always the same ($T > 0$), and permutation tests on the size of significant clusters showed that both samples

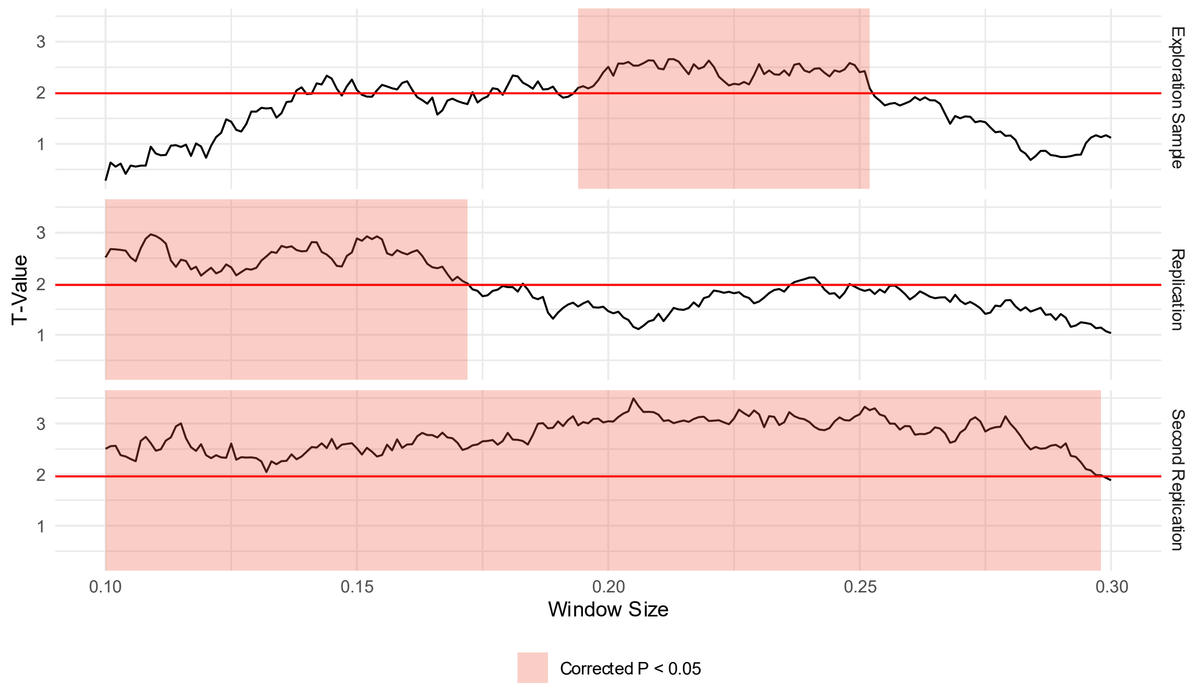


Supplementary Figure 4. Window size analysis reveals significantly larger clusters of above threshold effects than would be expected under the hypothesis that upward and downward trials are exchangeable. The window size corresponds to the absolute distance from the center of the screen in which data is included (i.e., a window size of 0.25 stretches from 25% to 75% of the screen height). The black line shows the T-Value of the t-test against zero of participant correlation coefficients between the distance the ball had moved at the time of flash and the overshoot. As the window size increases, more data is included but the edge effects caused by screen size are more prominent. Shaded regions show significantly large clusters of the T-Value being above the P<0.05 threshold.

have areas that are statistically larger than would be expected if no effect was present. Therefore, in the Second Replication sample, we pre-registered this permutation test to determine if a repulsive effect was detectable through varying the size of the analysed window. In the second replication, as can be seen, the effect was significant across the entire window. These analyses give greater confidence in the presence of effect in Experiment 3, despite the effect not quite crossing threshold ($p=0.06$) in the prespecified window in one of the three studies.

#### 5. Exploration of Previously-Reported Opposite Effect

##### *5.1 Motivation*

A previous study employed a similar paradigm to that in Experiment 3 but objects would disappear at a certain point along the trajectory and require reproduction, rather than the reproduction involving the location of a flash (Hubbard, 2001). Hubbard (2001) interestingly reported that the overshoot was larger for objects vanishing at the bottom, relative to top, of the screen. This was interpreted as an attraction toward the expectation of gravity causing objects to accelerate downwards, relative to upwards, and in direct conflict with the results from Experiment 3. Therefore, we sought here to replicate the findings of Hubbard (2001) in a large online sample. For full details see the pre-registration: https://osf.io/q2etu. To pre-empt, across all analyses, we did not replicate Hubbard (2001).

##### *5.2 Results*

Specifically, we tested whether the height on the screen at the time of vanishing would affect the overshoot using a pre-registered Bayesian repeated measures analysis of variance (rmANOVA). A Bayes factor of 37.9 shows that there is an effect of height on participants' overshoots. This significant

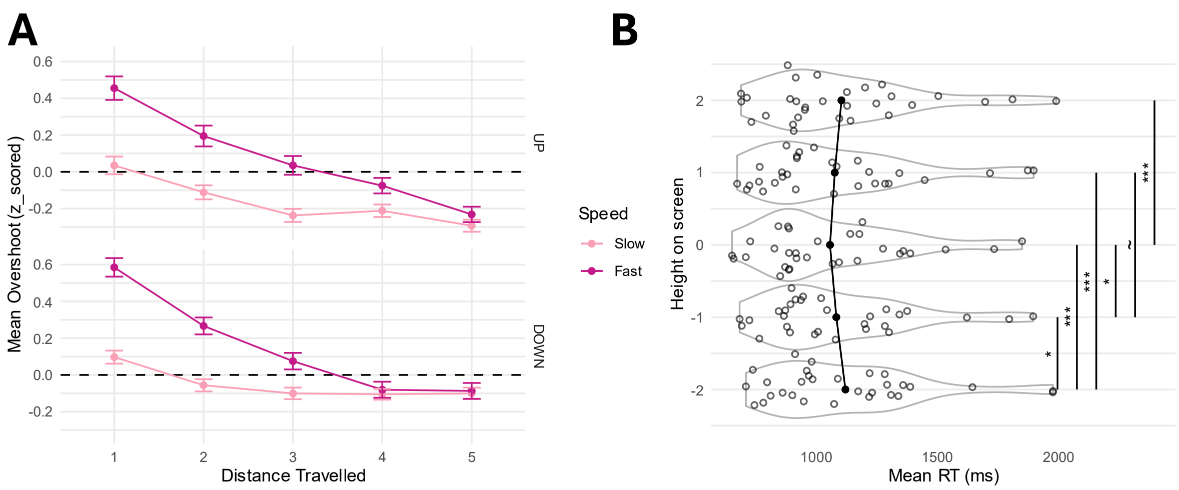


Supplementary Figure 5. **A** Participant overshoots as a function of the distance travelled on the screen. Fast and slow trials are separated with the group means plotted with error bars of the standard error of the mean. Participant overshoots were largest when objects had travelled the smallest distance, and these overshoots decreased with time, perhaps as participants accumulated more evidence. **B** Participant mean reaction time (x axis) or each disappearance height (y axis). Participant reactions were generally faster in the middle of the screen. Significance bars included if the pairwise Bayesian t-test had a $BF > 3$ or $BF < 1/3$ ($* = BF > 3$; $** = BF > 10$; $*** = BF > 30$; $\sim = BF < 1/3$).

effect stopped our data collection ($n = 30$) as a Bayesian stopping rule of $BF > 5$ or $BF < 0.2$ was pre-registered.

To parallel the original analysis in Hubbard (2001), we first performed a frequentist 5 (height) x 2 (direction) x 2 (speed) x 3 (size) rmANOVA. This ANOVA found a significant main effect of Height ($F(4{,}116) = 6.91, P < 0.001$) and Speed ($F(1{,}29) = 12.81, P < 0.001$). These main effects reflect that faster objects generated a larger overshoot, and that the smallest overshoot was in the middle of the screen, with larger overshoots at the top and bottom. Additionally, an interaction between Height and Direction ($F(4{,}116) = 11.00, P < 0.001$) showed that the larger overshoots occurred for upward motion at the bottom of the screen and downward motion at the top of the screen. Finally, the Height x Direction interaction also interacted with the Speed ($F(4{,}116) = 13.94, P < 0.001$) and the Size ($F(8{,}232) = 3.20, P < 0.01$) of the object. These three-way interactions show that the Height x Direction interaction is stronger for faster and larger objects.

Assuming linear relationships for effects of Height and Size, we can also transform the discrete identifiers used thus far into continuous variables, for better comparability with our own Experiment 3. When conducting a linear model analysis to examine the interactions between Height, Direction, Speed, and Size, with a random intercept for participant, the main effect of Height no longer persisted. Instead, a main effect of Direction was found ($\beta = -4.7 \times 10^{-2}; P = 1.2 \times 10^{-8}$), supporting previous location prior findings (De Sá Teixeira & Hecht, 2014; Hubbard, 1997). Additionally main effects of Speed ($\beta = 1.1 \times 10^{-1}; P < 2 \times 10^{-16}$) and Size ($\beta = 2.7 \times 10^{-2}; P = 7.8 \times 10^{-3}$) support previous findings in the representational momentum literature (Freyd & Finke, 1986). An interaction between Height and Direction ($\beta = -1.1 \times 10^{-1}; P < 2 \times 10^{-16}$) interacted with Speed ($\beta = -5.4 \times 10^{-2}; P < 2 \times 10^{-16}$) and Size ($\beta = -3.0 \times 10^{-2}; P = 3.2 \times 10^{-5}$), supporting the frequentist rmANOVA results.

We also tested whether the height on the screen at the time of vanishing affected the reaction times of participants using a pre-registered Bayesian rmANOVA. A Bayes factor of 66.0 provided strong evidence for an effect of height on participant's reaction times. Pairwise comparisons of mean reaction times at each vanishing height with Bayesian paired t-tests showed that balls vanishing in the middle of the screen (Mean RT = 1061ms) were responded to more quickly than those at the top (Mean RT = 1108ms) or bottom (Mean RT = 1125ms) of the screen (See Supplementary Figure 5B).

**6. Linear Modelling of Experiment 3**

Analysis of the data in all sub-experiments of Experiment 3 was performed with linear mixed effects modelling. We compared three models using the Akaike Information Criteria (AIC) to determine which best accounts for the data. The overshoot was predicted from the speed of the ball at the time of the flash, the direction of motion of the ball, and the distance that the ball has travelled. In the null model, only the speed and direction were modelled as these relate to the previously observed representational momentum and representational gravity (location prior) effects. An intermediary distance model included the distance travelled by the ball as a predictor of overshoot. Finally, the model of interest allowed an interaction between the distance travelled and the direction of motion. All three models included a random intercept and random slopes for the predictors for each participant.

$$null = lmer(Overshoot \sim Speed + Direction + (1 + Speed + Direction + Distance|PID))$$
$$distance = lmer(Overshoot \sim Speed + Direction + Distance + (1 + Speed + Direction + Distance|PID))$$
$$interaction = lmer(Overshoot \sim Speed + Direction * Distance + (1 + Speed + Direction + Distance|PID))$$

The interaction model ($AIC = -179{,}206.5$) had a more negative AIC than both the null ($AIC = -179{,}190.2$ ) and the distance model ($AIC = -179{,}177.0$). In the interaction model. The intercept estimate was positive ($\beta = 7.998 \times 10^{-10}$; $P < 0.001$) and reflects a general tendency to overshoot, which was larger when the ball was travelling faster ($\beta = 2.246 \times 10^{-3}$; $P < 0.001$) - consistent with representational momentum (Freyd & Finke, 1986). Evidence for the attractive effect of location priors (De Sá Teixeira & Hecht, 2014; Hubbard, 1997) was found such that downward moving balls had a larger overshoot than upward moving balls ($\beta = -9.357 \times 10^{-3}$; $P < 0.001$). The distance travelled did not significantly predict the overshoot ($\beta = -7.457 \times 10^{-4}$; $P = 0.519$) but the interaction between distance and direction was significant ($\beta = 2.208 \times 10^{-3}$; $P < 0.001$). This final interaction shows that downward moving balls had a larger overshoot at the *start* of motion than they did at the end, whereas upward moving balls had a larger overshoot at the *end* of motion than they did at the start. Thus, this analysis supports the results in Experiment 3 that overshoots tend to be larger at the top of the screen than the bottom.
